# Genomics highlight an underestimation of the urban heat island effect on red oak phenology

**DOI:** 10.1101/2022.08.03.502691

**Authors:** M. Blumstein, S. Webster, R. Hopkins, D. Basler, D. L Des Marais

## Abstract

The phenological timing of leaf-out in temperate forests is a critical transition point each year, altering the global climate system via changes to carbon and hydrologic cycles and land-surface albedo. In turn, climate is impacting phenology by advancing leaf-out an average of 2.8 +/-0.35 days per decade as the planet warms. Thus, understanding the relationship between warming and leaf-out is critical for understanding future global change. Rural-to-urban gradients in temperature, which result in clines over which urban areas are up to 4°C warmer than their surrounding countryside (dubbed the urban heat island – UHI – effect), may be used as a space-for-time substitution in studies of response to climate change. However, studies have recently highlighted that using the UHI effect over space rather than measurements at the same site through time results in consistently weaker relationships between temperature and leaf-out date in spring (i.e., urban-to-rural gradients underpredict the impact of warming on leaf-out). While such studies suggest many potential environmental explanations, the effect of genetic diversity is often neglected. While sensitive to environmental warming, leaf-out phenology is also highly heritable. Given that rural areas are largely natural sites, they likely have higher intraspecific genetic diversity than urban sites, where plants are selected by land managers for a narrow set of resilience traits. Here we measured the environment, genomic background, and phenological timing of northern red oak (*Q. rubra*) over several years between an urban and rural site to demonstrate how genetic background explains why the UHI effect leads to an underprediction of plant response to warming. Using a space-for-time substitution, we found that the date of leaf-out at our sites is predicted to advance approximately 20 days over the next 80 years. However, if we further account for the genomic background at the two sites, leaf-out, phenology is predicted to advance 22 days; a 10% difference between the two models. We demonstrate that this stronger relationship is because urban trees are largely a monoculture and, moreover, are most closely related to individuals at the rural site that leaf out latest. We highlight the critical need to consider genetic background, particularly in studies examining highly heritable traits, because both environment *and* genetics are changing across rural-to-urban gradients.

## Introduction

Phenology, or the timing of recurring life events in organisms, is a critical adaptive trait that is being altered by climate change (Parmesan & Yohe, 2003; Visser & Both, 2005; Cook *et al*., 2012). Over the past century, phenological timing of plants has advanced 2.8 +/-0.35 days/decade in response to global warming (IPCC, 2014). The earlier onset of spring creates feedbacks with the climate system, as spring leaf-out is both advanced by warming temperatures and because the appearance of leaves alters global CO_2_/H_2_O fluxes and the land surface albedo (Schwartz *et al*., 2006; Penuelas *et al*., 2009; Richardson *et al*., 2013).

While the climate is warming globally, urban areas are on a steeper trajectory than their surrounding rural areas, a phenomenon referred to as the urban heat island (UHI) effect (EPA, 2022). Ambient temperatures in cities are elevated relative to the surrounding countryside by up to 4°C during the day and by 3°C at night, primarily due to surface radiation from unvegetated, paved landscape (eg. EPA, 2022). As a result, UHI’s are often used as space-for-time substitutions to predict how warming temperatures will impact the phenology of species in a region (eg. Melaas *et al*., 2016; Zhao *et al*., 2016; Wang *et al*., 2019). For example, studies utilizing Landsat data have found urban trees in the Midwestern and Northeastern united states have up to a 15 day (Zipper *et al*., 2016) or 18-22 day (Melaas *et al*., 2016) increase in growing season length, respectively, compared to trees in nearby countryside.

More recently however, doubt has been cast on the validity of using rural-to-urban gradients as space-for-time substitutes (eg. Wohlfahrt *et al*., 2019). Studies have highlighted that using rural-to-urban gradients underestimates phenological advances in leaf out timing (Wohlfahrt *et al*., 2019) and the strength or relationship between leaf out timing and temperature (Li *et al*., 2019; Meng *et al*., 2020) when compared to measures taken at the same sites over time. While these studies examined several potential confounding environmental explanations for their findings, they neglected the possible role that within-species genetic diversity plays in shaping these patterns. This omission is potentially large given phenological timing of leaf out is a highly heritable trait, with broad-sense heritability (H^2^) estimates ranging between 0.6-0.9 for tree species (Yao & Mehlenbacher, 2000; Alberto *et al*., 2011; McKown *et al*., 2014). Furthermore, this heritable variation often shows strong patterns of local adaptation (eg.Hall *et al*., 2007; Alberto *et al*., 2011; Evans *et al*., 2014), indicating that variation in phenological timing is not only plastic, but also strongly governed by genetics.

Examining the same sites, and consequently the same genotypes, through time is a direct measure of plasticity. For a space-for-time substitution to be a comparable measure of plastic variation, we must assume that genetic variation is constant across that gradient. However, while rural areas represent natural populations, urban environments are managed, effectively representing artificial selection favoring traits that provide resiliency to urban stressors. As a result of this artificial selection, we predict that urban tree populations have reduced genetic variation compared to rural ones. Here we demonstrate how the inclusion of genomic differences can explain why rural-to-urban gradients are a poor space-for-time substitution for phenological timing using the model species Northern Red Oak (*Q. rubra*) (Figure 1). We demonstrate the importance of accounting for the often-overlooked heritability of phenological timing in trees in efforts to understand the impacts of global change on leaf out.

**Figure 1.**
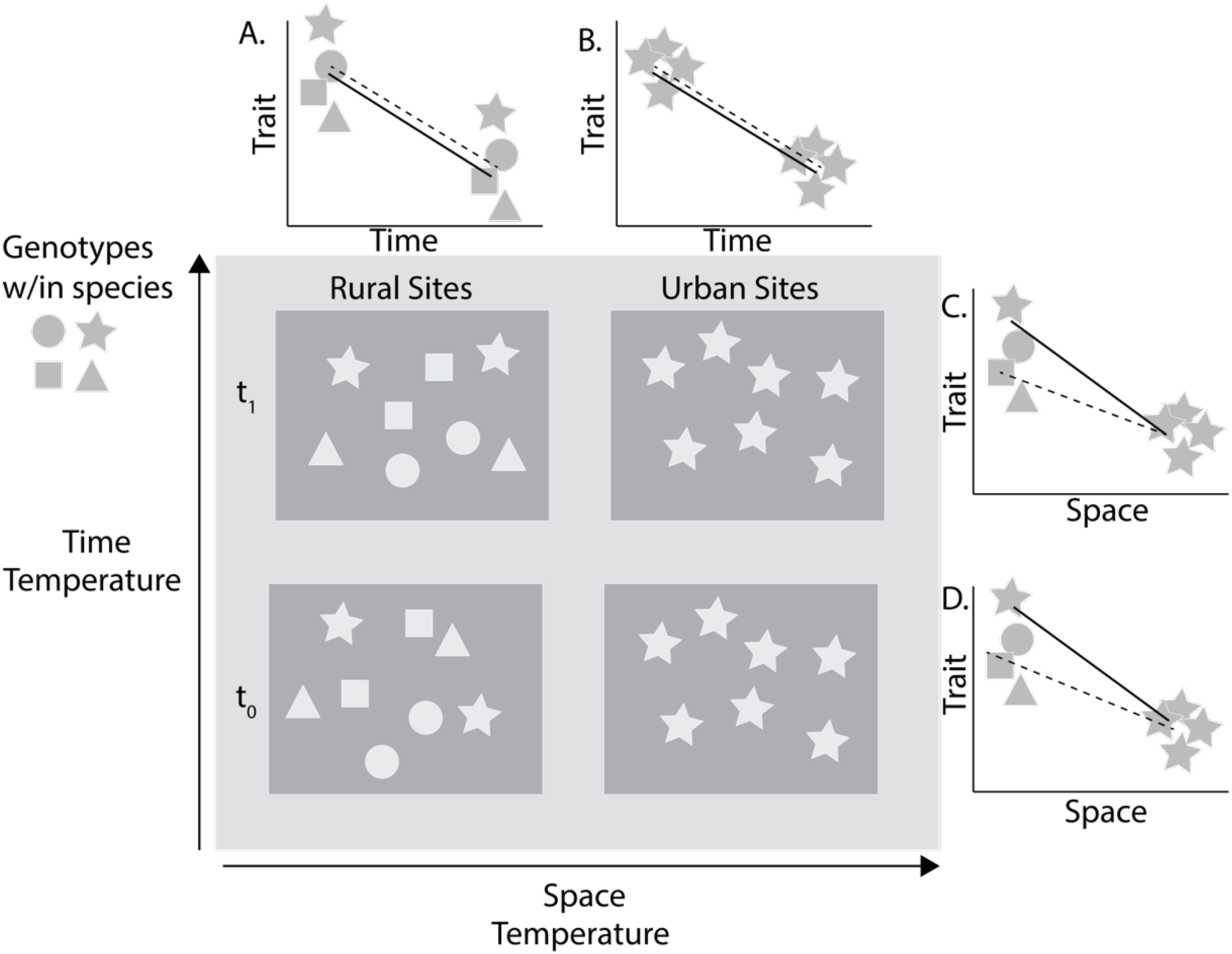
A diagram illustrating how accounting for differences in genetic diversity across sites can alter predictions. Rural sites (left) have high within species genetic diversity, while urban sites (right) have low diversity. When comparing the same site through time (A-B), genetic background remains the same and only temperature changes. Thus, the slope of the line accounting for genetic differences & environment (solid line) is the same as the one for just environment (dashed line). However, in this scenario, when comparing different sites across a temperature gradient (C-D), the slope of the line accounting for genetics & environment (solid) is much steeper than that accounting for just environment (dashed). Thus, if there are differences between two sites in genetic background and the trait is heritable (ie. trait means differ by genotype), then model predictions will be off when compared to the same sites through time.

## Methods

### Site Description

This study was conducted in the city of Cambridge, Massachusetts, USA (Urban Site; 42.3784º N, −71.1479º W) and in the Prospect Hill tract of the Harvard Forest, a well-described research forest in Petersham, Massachusetts (Rural Site; 42.5377º N, 72.1715º W). Petersham has a population just over 1,000, annual average temperatures of 7.97 ºC and 110 cm of precipitation (National Climatic Data Center, 2021; Boose and Gould, 2022). Cambridge is a small metropolis with just over 100,000 people and is the 9^th^ most densely populated urban center in the United States in 2020 with 29 people/acre (US census). Cambridge has a mean annual temperature of 11.4 ºC and mean average precipitation of 110 cm.

Within Petersham, Harvard Forest is a 3,000-acre area dedicated to ecological research. It is a mature, temperate forest with some woody wetlands, populated mainly by Northern Red Oak (*Quercus rubra*), Red Maple (*Acer rubrum*), and Hemlock (*Tsuga canadensis*) (Smith et al., 2019; Klosterman and Richardson, 2017). The Prospect Hill tract contains within it a ForestGEO (Global Earth Observatory Network) plot (https://forestgeo.si.edu/), a 35-hectare region with every stem above 1 cm mapped and measured for DBH (SI geo dataset at HF) on five year intervals. All measurements taken at the Harvard Forest occurred within this ForestGEO plot. Notably, the plot also contains a 3.5-acre swamp with standing water year-round. Aerial drone surveys (described below) and genomic sampling fell along a gradient within the ForestGEO plot from the swamp to drier areas (Figure S1A).

Cambridge has over 19,000 trees, featuring 93 unique species among its street tree population; most abundant are Norway Maple (*Acer platanoides*), Honeylocust (*Gleditsia triacanthos*), and Red Maple (*Acer rubrum*). Red Oak comprise 2.2% of the city-owned street tree population in Cambridge (Boukili, 2016).

### Phenological Observations

We selected 15 trees around Harvard Square in Cambridge and 43 trees in Harvard Forest to represent urban and rural red oak populations. Phenological observations made at the Harvard Forest were via unmanned aerial vehicle (UAV) photography during the growing seasons in 2014, 2015 (Klosterman *et al*., 2017) and 2018 (Basler, unpublished). Focal trees were chosen for study (N = 43) across a microenvironmental gradient from the standing swamp to a drier portion of the ForestGEO plot, representing a diverse phenotypic spread. Individual trees were identified by overlaying the georeferenced ForestGEO stems on the aerial imagery and then drawing tree crowns by hand. The green-chromatic coordinates or GCC were derived from these images for each individual using R and the phenopix package v.2.4.2. Seasonal transition dates were found by fitting a Beck spline to the data and estimating derivative transition points. Start of season (SOS) is one of the metrics derived from this process and used here as leaf-out date.

Cambridge trees span a gradient from the Charles River to inner Cambridge and lie in three different jurisdictions (Harvard University, The City of Cambridge, and the Charles River Conservancy; N = 30). Phenology observations were made via human observation using a modified version of the BBCH scale to capture major transition points for *Quercus rubra*. Data were collected in the spring of 2020 and 2021. Our stage 3, when leaves are visible but have not escaped the bud yet, corresponds to SOS found via aerial images.

### Genomic Analyses

Leaf tissues were collected in May just after budburst via shotgun at the Harvard Forest and pole pruner in Cambridge. Samples were prepared with Qiagen DNeasy kits, following their standard protocols. Low coverage whole-genome sequencing (lcWGS) was performed at the MIT BioMicro Center. The DNA library was prepared by the BioMicro Center using Nextera Flex and sequencing was performed on an Illumina NovaSeq.

To align sequence reads to the *Q. rubra* reference genome (https://phytozome-next.jgi.doe.gov/info/Qrubra_v2_1), we indexed the reference genome and aligned the sequences to the assembly using bwa and generated binary alignment map (BAM) files using SAMtools (Li, 2013; Li et al., 2009). We called SNPs using freebayes, creating a master VCF (variant call format) file (Garrison and Marth, 2012). Using PLINK, we pruned the VCF for linkage disequilibrium, filtered for minor allele frequency greater than 0.1, and specified biallelic sites before calculating the eigenmodes of the SNP marker data matrix (Purcell et al., 2007). All bioinformatics tools were used with default settings. To establish relatedness amongst our samples, we utilized the first two eigenvectors derived from our SNP matrix.

### Statistical Analyses

We compared the growing degree days (GDD) on May 1 at each site in each year of observation. GDDs were calculated using daymet data and the daymetr and pollen packages in R. The temperature base was set to 5°C. We used linear models in R, with day-of-year (DOY) of leaf-out as our dependent variable. Our null model fit GDD as a predictor, while our genomic model fit PC1and PC2 in addition to GDD, both of which were derived from our SNP matrix (Figure 2).

**Figure 2.**
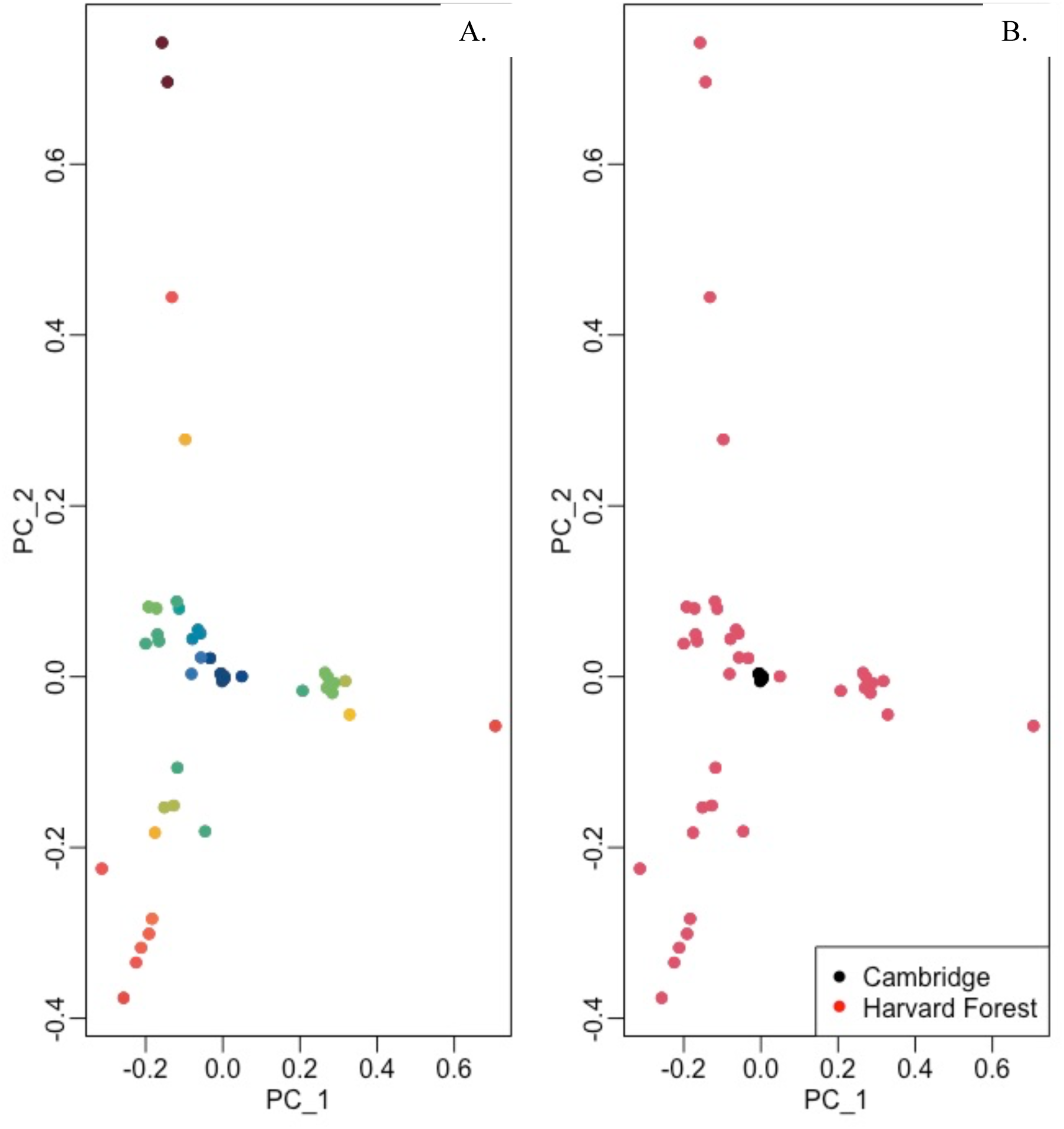
PCA of SNP matrix (N_SNPs_ = 591,818, m.a.f > 0.1). Panel (A) shows samples colored by genotype index – the absolute value of the product of PC1 by PC2. Panel (B) shows samples colored by site they were collected at.

## Results

### Genomic Analyses

Across our 58 sequenced individuals, genomes yielded 85% coverage and 5x depth across all linkage groups and scaffolds (12 LG in *Q. rubra*). After all quality control steps were taken, we retained a set of 58 individuals and 591,818 SNPs. A plot of the first and second PCs shows clustering of the Cambridge oaks in the center-left, suggesting very little genetic diversity among the urban trees (Figure 2). While overlapping with the urban trees, Harvard Forest oaks are more widely spread throughout the PC space, indicating extensive genetic variation throughout the study area (Figure 2).

### Phenological Analyses

On average, Harvard Forest Red Oaks leafed out on julian day 138 (s.d. = 4.7) across all years and Cambridge trees leafed out on julian day 128 (s.d. = 9.3).

### Phenotype = Genotype + Environment

Our model running the DOY of budburst versus the number of GDDs on May 1, at each site/year combination was significant, with a negative slope indicating leaves come out earlier as temperatures rise in spring (m = −0.07, R^2^ = 0.39, p < 0.001; Figure 3). Our model of DOY of budburst versus the number of GDDs on May 1 and genotype, as given by PC1 and PC2 was also significant, with more of the variance explained and a more negative slope by adding genotype (m = −0.09, R^2^ = 0.56, p < 0.001; Figure 3).

**Figure 3.**
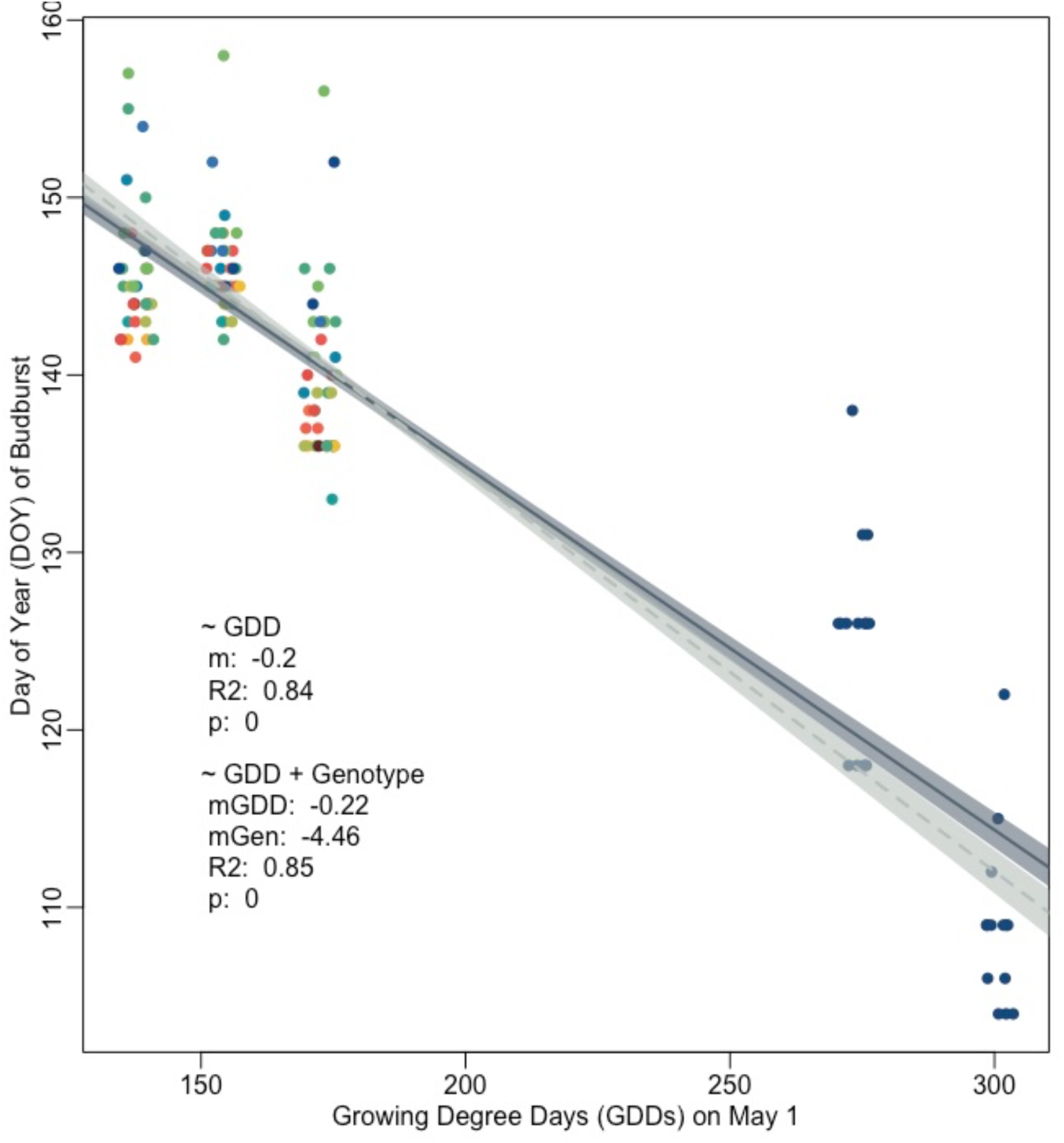
The day of year (DOY) of budburst of individual red oaks growing at the Harvard Forest (rural site) and in Cambridge (urban site) versus the growing degree days (GDD) on May 1 of that year. Three years of data for the Harvard Forest (2014, 2015, & 2018) and 2 years of data for Cambridge (2020, 2021) are shown. The solid line shows a linear model fit using only environment (GDD on May 1) as a predictor, with the standard error shaded in dark blue-gray. The dashed line shows a linear model that additionally includes the genotype of each individual as described by the first two PCs of their underlying SNP matrix, with standard error shaded in light gray. Dots are jittered to show more of individuals sampled and colored by genotype (absolute value of PC1 x PC2).

## Discussion

We demonstrate that rural-to-urban gradients in temperature underpredict the sensitivity of leaf-out in trees to warming when genetic variation is not included in the model. We found individuals of our species growing in urban areas are very genetically similar (Figure 2B). This makes as urban trees are usually sourced from a small number of nurseries and are chosen for very specific traits that enable resiliency in the urban environment. Conversely, our wild population of *Q. rubra* is wind pollinated and occupies a diversity of microclimates, leading to a high degree of genetic diversity at any given site (Cavender-Bares, 2019). As a result, not accounting for these differences between sites leads to a 3-day or 29% difference in predicted leaf-out date given warming temperatures over the next 80 years.

Urban tree managers tend to select cultivars from species that are tolerant of extreme conditions or that have a nice presentation (McElhinney & Harper, 2019). Given that leaf-out phenology tends to be highly heritable in trees (eg. Yao & Mehlenbacher, 2000; Alberto *et al*., 2011; McKown *et al*., 2014), by selecting only a few cultivars or genotypes to plant out, city managers are inadvertently biasing the leaf-out date of each species growing in the urban environment. This is the case for northern red oak growing within Cambridge. While the street trees in or sample span three different management entities (Harvard University, Charles River Conservancy, & The City of Cambridge), they are all clustered together indicating very little underlying genetic diversity (Figure 2B). This cluster of trees is most closely related to those that tend to leaf out on the later side at the Harvard Forest, our rural site (Figure 4).

**Figure 4.**
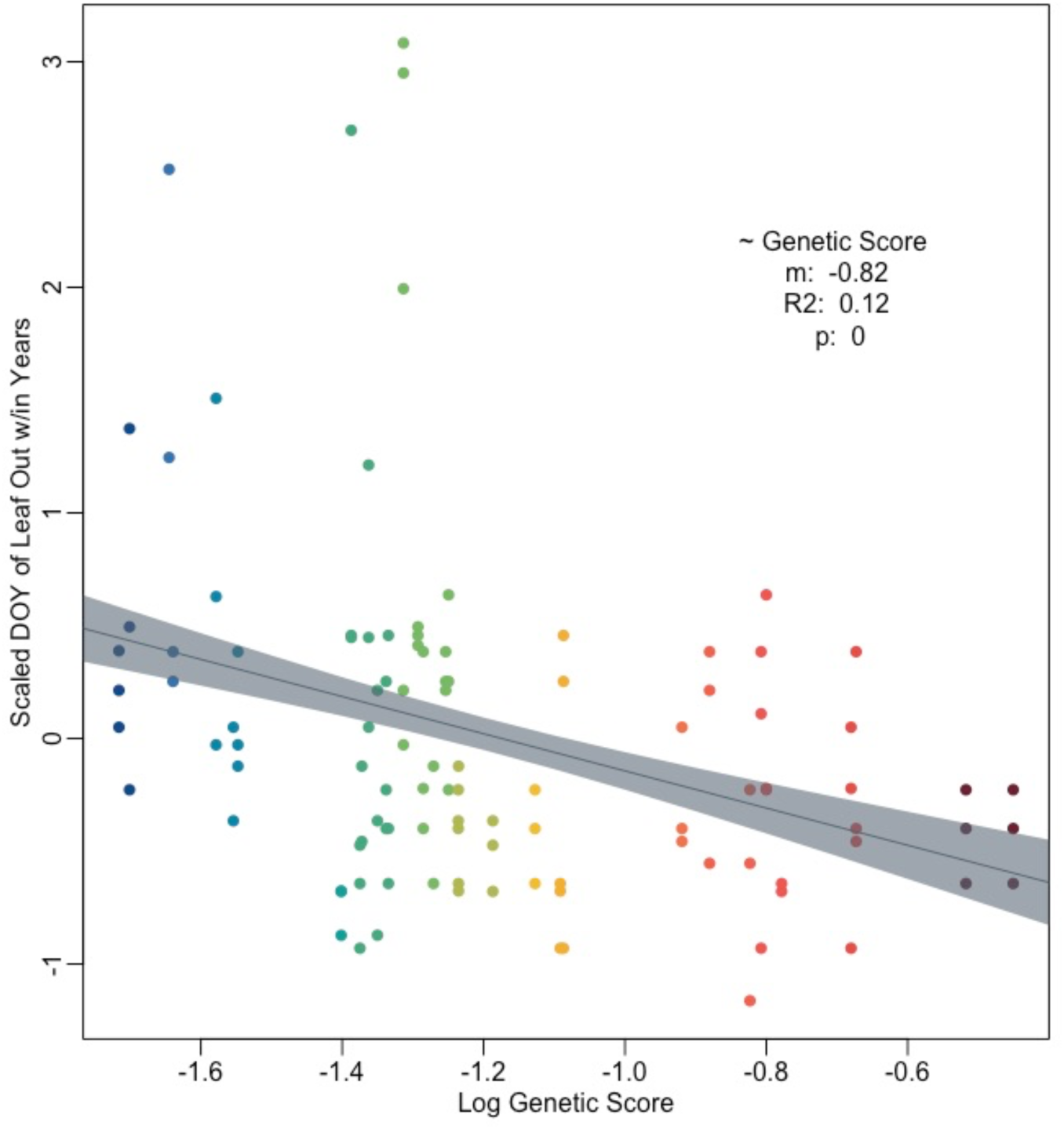
Day of year of leaf-out at the Harvard Forest across 2014, 2015, and 2018, by genotype -- as defined by the absolute value of the product of PC1 and PC2. The results of each linear model comparison is presented in text. Points are colored by genotype.

While striking, our data does come with some caveats. We did not control for microenvironmental variation at either site, which may impact leaf-out times. However, microenvironmental variation would likely only add noise to our data and dappen our trend, rather than enhance it. Another potential concern is the impact of ontogeny on phenology, as younger trees can leaf out earlier than their elder counterparts (Vitasse, 2013). As urban trees tend to have shorter life spans than rural trees (eg. Smith *et al*., 2019), the average age of our trees is likely younger in our urban setting. However, we only sampled from the largest size class at both sites, so while not the same age, our individuals were large canopy trees and thus the ontogenetic effects that have been cited when comparing seedlings or saplings to canopy trees have little affect our study (eg. Vitasse, 2013). Finally, we took great strides to account for our different methods of making phenological observations at the rural (aerial drone) versus urban sites. We used ground observations taken at the Harvard Forest to ensure our on the ground start-of-season metric matched those derived from aerial images (Figure S1;Klosterman & Richardson, 2017). We additionally ensured that our across site differences were aligned with external metrics using phenocams pointed at red oaks at both sites (Figure S2;Richardson *et al*., 2018).

Estimating the timing of leaf out in temperate trees is a critical transition point in dynamic vegetation models, where small changes can have large consequences for our warming predictions. We show not only that using the UHI effect can underestimate warming, but we demonstrate for the first time why, providing an explanation for a trend highlighted in previous studies (Wohlfahrt *et al*., 2019; Meng *et al*., 2020). Our findings add to a literature highlighting the importance of considering genetics in ecological studies and provides a path forward for once again using rural-to-urban gradients as large-scale experiments.

Our findings also provide important information for urban street tree management. While the climate is warming globally, cities in particular are experiencing unprecedented heat waves. Trees have been cited as a way of mitigating not only the global impacts of warming, but also combatting the UHI effect on a local level. Beyond their innumerate health and social benefits, transpiration and shade from trees can greatly cool neighborhoods (Turner-Skoff & Cavender, 2019). For example, the combination of evapotranspiration and shading in trees can reduce peak summer temperatures by 1-5°C (EPA, 2022). A longer growing season could lead to a greater offset of the UHI in a city and health benefits for its citizens.

Our results also aid land managers in rural areas seeking to undertake assisted migration, or the movement of genotypes northward to speed up adaptation to climate change. We’ve identified high genetic diversity at the Harvard forest and a large spread in leaf-out date, a locally adapted trait in trees that enables success in any given population. Genes that enable trees to leaf-out earlier will be critical under a warming climate to keep species competitive. We identify not only a range of genotypes, but also a range of microenvironments that they tend to be found in related to soil moisture (Figure S2). While more study is needed, our results point to the promise of mapping microenvironmental variation to genetic patterns for large-scale identification of candidates for assisted migration.

## Author Contributions

Conceptualization: MB, SW, RH, DDM

Methodology: MB, SW, DB

Funding Acquisition: MB, SW, & DDM

Visualization: MB & SW

Writing – original draft: MB, SW, & DDM

Writing – review & editing: MB, SW, DB, DDM, RH

## Data and materials availability

All data are available in the main text or the supplementary materials

## Supplementary Materials

